# A SCID mouse model to evaluate the efficacy of antivirals against SARS-CoV-2 infection

**DOI:** 10.1101/2022.05.13.491916

**Authors:** Rana Abdelnabi, Caroline S. Foo, Suzanne J. F. Kaptein, Robbert Boudewijns, Laura Vangeel, Steven De Jonghe, Dirk Jochamns, Birgit Weynand, Johan Neyts

## Abstract

Ancestral SARS-CoV-2 lacks the intrinsic ability to bind to the mouse ACE2 receptor and therefore establishment of SARS-CoV-2 mouse models has been limited to the use of mouse-adapted viruses or genetically modified mice. Interestingly, some of the variants of concern, such as the beta B.1.351 variant, show an improved binding to the mouse receptor and hence better replication in different Wild type (WT) mice species. Here, we desribe the establishment of SARS-CoV-2 beta B.1.351 variant infection model in male SCID mice as a tool to assess the antiviral efficacy of potential SARS-CoV-2 small molecule inhibitors. Intranasal infection of male SCID mice with 10^5^ TCID50 of the beta B.1.351 variant resulted in high viral loads in the lungs and moderate signs of lung pathology on day 3 post-infection (pi). Treatment of infected mice with the antiviral drugs Molnupiravir (200 mg/kg, BID) or Nirmatrelvir (300 mg/kg, BID) for 3 consecutive days significantly reduced the infectious virus titers in the lungs by 1.9 and 3.8 log10 TCID50/mg tissue, respectively and significantly improved lung pathology. Together, these data demonstrate the validity of this SCID mice/beta B.1.351 variant infection model as a convenient preclinical model for assessment of potential activity of antivirals against SARS-CoV-2.

**Importance:** Unlike the ancestral SARS-CoV-2 strain, the beta (B.1.351) VoC has been reported to replicate to some extent in WT mice (species C57BL/6 and BALB/c). We here demonstrate that infection of SCID mice with SARS-CoV-2 beta variant results in high viral loads in the lungs on day 3 post-infection (pi). Treatment of infected mice with the antiviral drugs Molnupiravir or Nirmatrelvir for 3 consecutive days markedly reduced the infectious virus titers in the lungs and improved lung pathology. The advantages of using this mouse model over the standard hamster infection models to assess the *in vivo* efficacy of small molecule antiviral drugs are (i) the use of a clinical isolate without the need to use mouse-adapted strains or genetically modified animals (ii) lower amount of the test drug is needed and (ii) more convenient housing conditions compared to bigger rodents such as hamsters.

## Introduction

Since its emergence in China end of 2019, the severe acute respiratory syndrome coronavirus (SARS-CoV-2) has resulted in a global pandemic with officially >517 million cases (as of May 10, 2022) and around 15 million deaths as estimated by WHO (1). Several SARS-CoV-2 variants of concern (VoC), that result in immune escape and/or enhanced viral transmission have since then emerged (2, 3). Small animal models are necessary to study the virus-induced pathogenesis as well as to serve as preclinical tool to assess the efficacy of vaccine and therapeutics against the viral infection. Similar to SARS-CoV, SARS-CoV-2 enters the host cells through attachment to the cellular angiotensin-converting enzyme 2 (ACE2) (4). Since SARS-CoV-2 binds effeciently to the hamster ACE2 (5), Syrian hamsters are considered so far one of the best small animal models avaliable for SARS-CoV-2. On the other hand, the spike of the ancestral SARS-CoV-2 lacks the intrinsic ability to efficiently bind to the murine ACE2 (5) and hence this strain has a limited replication in WT mice. Consequently, alternative strategies have been developed to allow the establishment of mouse models for SARS-CoV-2. One of these strategies is adaptation of the virus in murine lung tissues to enhance the binding capacity to the murine ACE2 (6, 7). Other strategies focused on introduction of human ACE2 in wild-type mice either by transduction adenovirus or adeno-associated virus that expresses human ACE2 (8) or using genetically modified human ACE2 transgenic (9) or humanized mice (10). Unlike the ancestral strain, some of the evolved SARS-CoV-2 VoCs proved to carry spike protein mutations, mainly the N501Y, that enable efficient binding to the murine ACE2 and hence better replication in WT mice (11, 12). Besides the N501Y mutation, the spike of the beta B.1.135 variant carries the K417N mutation that was previously reported in a virulent mouse-adapted SARS-CoV-2 variant (13). Several studies have shown the ability of the beta variant to replicate to some extent in WT mice species such as C57BL/6 (11, 12) and BALB/c (14, 15). Here, we wanted to explore whether the beta SARS-CoV-2 variant replicates more efficiently in severe combined immune deficient (SCID) mice than in the reported wild type mice and whether in such case, SCID mice can be used to develop a suficienlty robust infection model to study the efficacy of small molecules inhibitors of SARS-CoV-2 infection.

## Results

First, a small pilot study was performed to assess the efficiency of replication of the beta (B.1.351) SARS-CoV-2 variant in male SCID mice in comparison to replication in immunocompetent male BALB/c and C57BL/6 mice. All mice (n=9 per species), were infected with 10^5^ TCID50 of the beta variant. At day 3 post-infection (pi), all animals were euthanized and lungs were collected to quantify the infectious virus titers. As expected, the infectious virus titer in the lungs of infected SCID mice (median TCID50/mg lung of 3.28×10^4^) was significantly higher than that observed in the lungs of infected BALB/c mice (median TCID50/mg lung of 3.26×10^3^, p=0.047) and C57BL/6 mice (median TCID50/mg lung of 2.05×10^3^, p=0.0054), **supplementary Figure S1**.

Next, we explored the kinetics of replication of the beta variant in SCID mice. To that end, 7-9 weeks old male SCID mice were infected intranasally with with 10^5^ TCID50 of the beta variant. At days 3 through 7 post-infection (pi), 10 animals were euthanized and lungs were collected to quantify the infectious virus titers. The highest infectious virus titers were observed at day 3 pi (**Fig. 1A**). From day 4 pi onwards, the infectious virus titers in the lungs were significantly lower than those observed at day 3 pi (**Fig. 1A**). A minor weight loss was observed on day 3 pi (average %body weight change of -0.8) after which animals started to gain weight normally (average %body weight change of 4 on day 4 pi) (**Fig. 1B**). When a group of 5 infected mice were monitored up to 14 days pi, no weight loss or any signs of morbidity were observed in this group (average %body weight change of 13 on day 14 pi). Histological examination of the lungs from infected mice at day 3 pi revealed mild signs of peri-bronchial inflammation, significant peri-vascular inflammation and intra-alveolar hemorrhage (**Fig. 1C**) with median cumulative pathology score of (4.5).

**Fig. 1.**
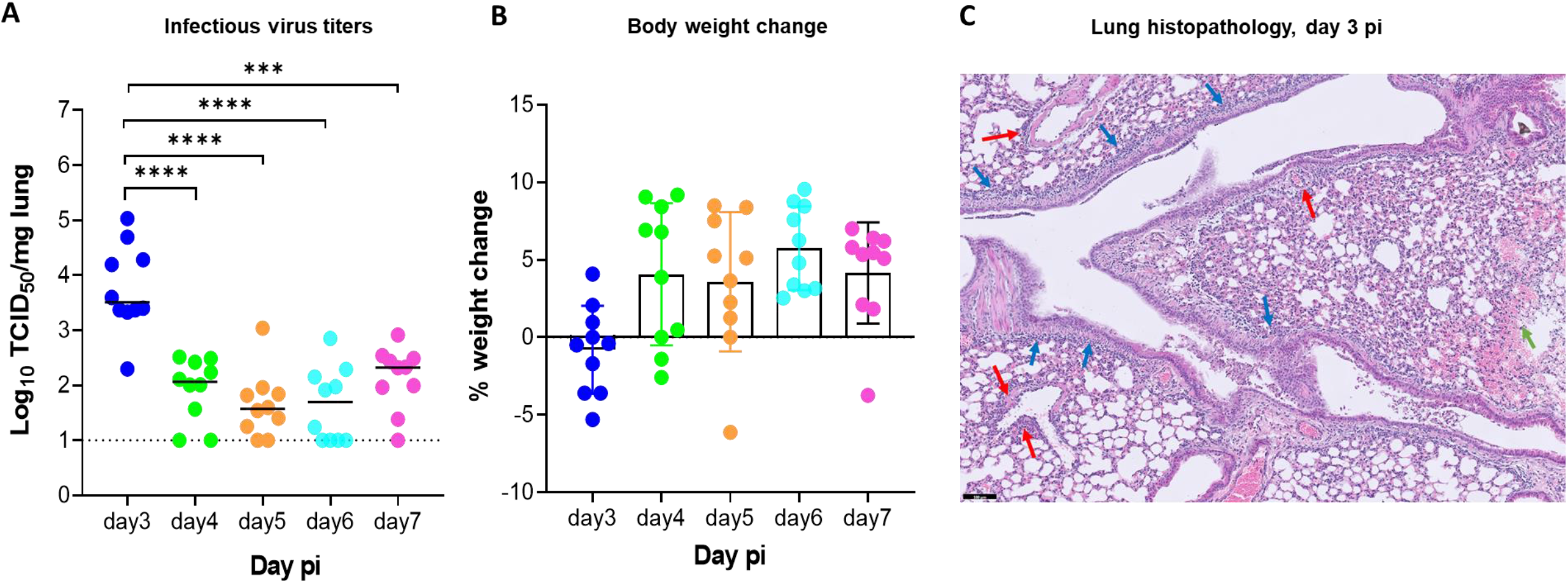
Replication kinetics of beta (B.1.351) SARS-CoV-2 in male SCID mice. (A) Infectious viral loads in the lungs of male SCID mice infected with 10^5^ TCID_50_ of beta SARS-CoV-2 variants at different days post-infection (pi) are expressed as log_10_ TCID_50_ per mg lung tissue. Individual data and median values are presented. Data were analyzed with the Mann™Whitney U test. ***P =0.0003, ****P < 0.0001 (B) Weight change at different days pi in percentage, normalized to the body weight at the time of infection. Bars represent means ± SD. All data are from 2 independent experiments with 10 animals per group. (C) Representative H&E image of lung from SCID mouse infected with the beta variant at day 3 pi showing limited peri-bronchial inflammation (blue arrows), significant peri-vascular inflammation (red arrows) and intra-alveolar hemorhage (green arrow). Scale Bar=100 µM

In case the infectious virus detected at day 3 post infection represents actively replicating virus, it should be possible to suppress replication by treating the animals with antiviral drugs. We therefore assessed the potential antiviral efficacy of two clinically relevant SARS-CoV-2 inhibitors i.e. Molnupiravir (EIDD-2801) and Nirmatrelvir (PF-332) against beta variant replication in SCID mice. Briefly, male SCID mice were treated twice daily by oral gavage with either vehicle, Molnupiravir (200 mg/kg) or Nirmatrelvir (300 mg/kg) for three consecutive days starting from the day of infection with the beta variant (**Fig. 2A**). Mice were euthanized at day three pi for collection of lung tissues. A significant reduction of viral RNA loads was observed in the Molnupiravir (0.8 log10 genome copies/mg tissue, p=0.0025) and Nirmatrelvir (2.8 log10 genome copies/mg tissue, p<0.0001)-treated groups as compared to the vehicle control (**Fig. 2B**). Moreover, treatment of mice with Molnupiravir and Nirmatrelvir significantly reduced the infectious virus titers in the lungs by 1.9 (p<0.0001) and 3.8 (p<0.0001) log10 TCID50/mg tissue, respectively compared to the vehicle-treated group (**Fig. 2C**). No infectious virus titers were detected in four (out of 14) and eight (out of 14) animals in the Molnupiravir and Nirmatrelvir-treated groups, respectively (**Fig. 2C**). A significant improvement of lung histopathology scores was also observed in both the Molnupiravir (p=0.025) and Nirmatrelvir (p=0.0007)-treated groups (**Fig. 2D**). No significant weight loss or clinical signs of adverse effects were observed in the compounds-treated groups (**Fig. 2E**).

**Fig. 2.**
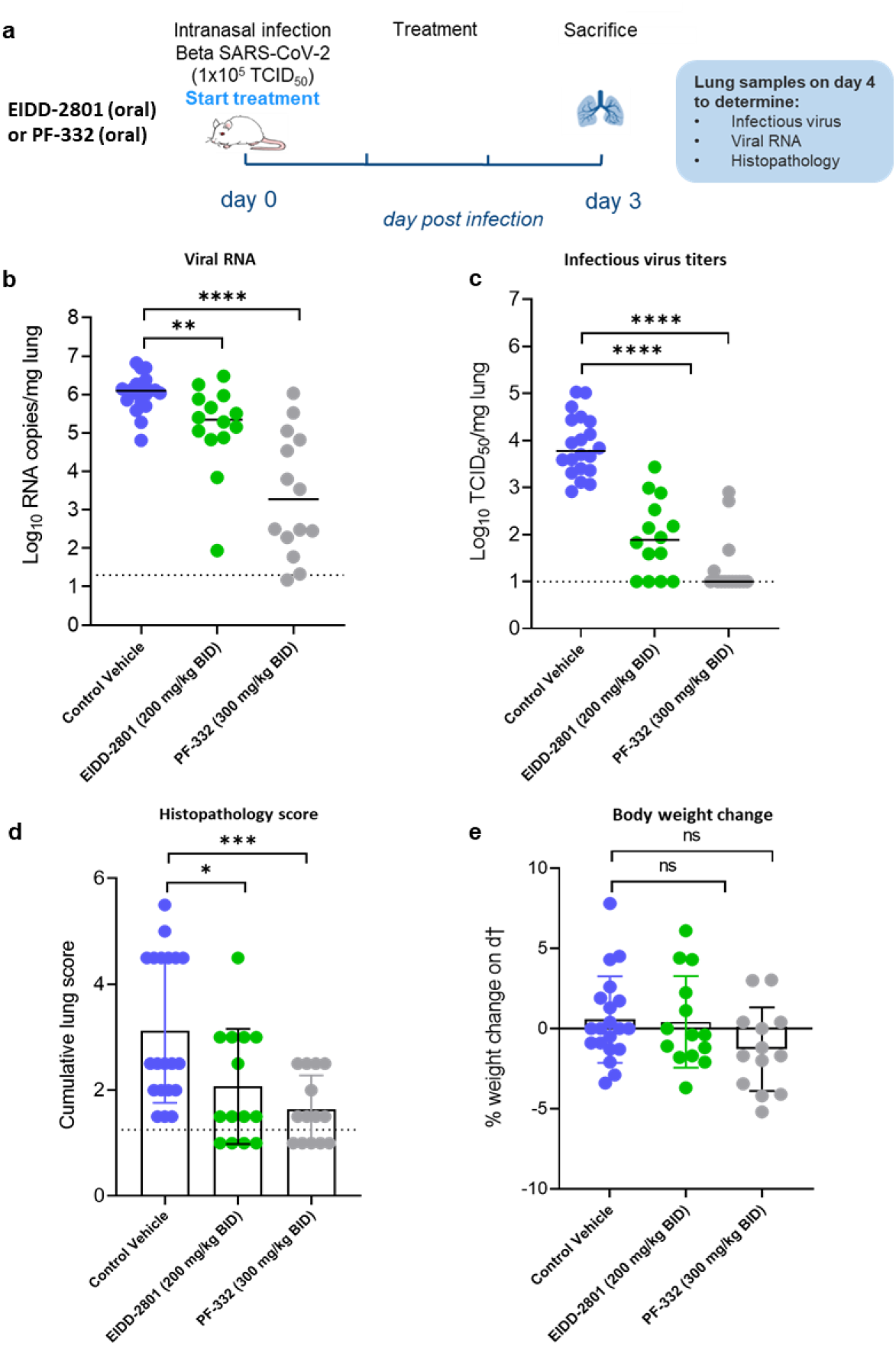
Molnupiravir (EIDD-2801) and Nirmatrelvir (PF-332) reduced viral loads in the lungs of beta (B.1.351) SARS-CoV-2-infected SCID mice. (A) Set-up of the study. (B) Viral RNA levels in the lungs of control (vehicle-treated), EIDD-2801-treated (200 mg/kg, BID) and PF-332-treated (300 mg/kg, BID) SARS-CoV-2 (B.1.351)™infected SCID mice at day 3 post-infection (pi) are expressed as log_10_ SARS-CoV-2 RNA copies per mg lung tissue. Individual data and median values are presented. (C) Infectious viral loads in the lungs of control (vehicle-treated), EIDD-2801-treated and PF-332-treated beta SARS-CoV-2™infected SCID mice at day 3 pi are expressed as log_10_ TCID_50_ per mg lung tissue. Individual data and median values are presented. (D) Cumulative severity score from H&E stained slides of lungs from control (vehicle-treated), EIDD-2801-treated (200 mg/kg, BID) and PF-332-treated (300 mg/kg, BID) SARS-CoV-2™infected SCID mice at day 3 pi. Individual data are presented and bars represent means ± SD. The dotted line represents the median score of untreated non-infected hamsters. (E) Weight change at day 3 pi in percentage, normalized to the body weight at the time of infection. Bars represent means ± SD. Data were analyzed with the Mann™Whitney U test. *P < 0.05, **P < 0.01, ***P < 0.001, ****P < 0.0001, ns=non-significant. All data (panels B, C, D) are from two independent experiments with 14 animals per group except for vehicle group (n=20).

## Discussion

The emergence of SARS-CoV-2 VoCs has raised a lot of concerns as these variants displayed the ability to escape vaccine-induced or naturally acquired immunity and to transmit faster than the ancestral strains. Besides, some of these variants have acquired certain mutations in the spike protein that allowed them to expand their host species (2, 12). The beta variant (B.1.351 or 501Y.V2) has been first reported in South Africa in October 2020 (16). The beta variant has acquired three mutations in the receptor binding domain (RBD) namely N501Y, K417N and E484K, in addition to other mutations in the spike and non-structural proteins (2). Among these mutations, the N501Y mutation (also presents in alpha variant) has been previously described in mouse-adapted viruses and proven to play an important role in increasing the affinity to the mouse ACE2 receptor (6). The K417N mutation has also previously been reported in a virulent mouse-adapted SARS-CoV-2 variant (13). In a pseudotype-based entry assay, the pseudoviruses carrying the beta variant spike attached more efficiently to the mouse ACE2 receptor than the alpha variants, suggesting that the K417N and E484K mutations in the RBD of beta variant may further enhance the binding to the mouse recptor (11). Recently, a comparative infection study in BALB/c mice revealed that the beta variant replicates more efficiently than the alpha and delta variant (15).

We here wanted to assess the infectivity of the beta SARS-CoV-2 variant in an immunodeficient mouse model i.e. SCID mice, with the aim to develop a robust SARS-CoV-2 mouse infection model for preclinical evaluation of potential antivirals. So far, the hamster SARS-CoV-2 infection model has been regarded as the best model to study the effect of antiviral agents, yet use of mice would facilitate such studies. We selected SCID mice as these animals are severely deficient in functional B and T lymphocytes and therefore they are believed to be more susceptible to viral infections than immunocompetent mice. Indeed, in our pilot infection study, the infectious virus titers of the beta variant in the lungs of infected SCID mice on d3 pi were significantly higher than that observed in the lungs of the immunocompetent BALB/c (1 log10 higher) and C57BL/6 (1.2 log10 higher) mice that were infected in parallel. Viral persistence in the lungs of SCID mice was observed in most of the infected animals up to 7 days pi. However, the infectious virus titers dropped significantly beyond day 3 pi. Therefore, day 3pi was selected as the endpoint for antiviral testing.

Nirmatrelvir (PF-332, Pfizer), is a potent inhibitor of the main protease Mpro (or 3CL protease) of SARS-CoV-2 and other coronaviruses (17). Paxlovid (Nirmatrelvir and Ritonavir tablets, co-packaged for oral use) have been authorized by FDA and EMA as well as by other regions. Molnupiravir (Lagevrio™, EIDD-2801, Merck) is the orally bioavailable prodrug of the ribonucleoside analogue N4-hydroxycytidine (NHC, EIDD-1931), which was initially developed for influenza (18) and has now also been approved by several countries/regions for the treatment of COVID-19.

We have previously shown that Molnupiravir (EIDD-2801) and Nirmatrelvir (PF-332) significantly inhibit the replication of the beta variant in Syrian hamsters (19, 20). Therefore, we used these two antiviral drugs to validate whether the SCID mice/beta variant infection model for antiviral studies. Treatment of beta variant-infected SCID mice for 3 consecutive days with Molnupiravir (200 mg/kg, BID) or Nirmatrelvir (300 mg/kg, BID) significantly reduced viral loads in the lung of infected mice with a potency close to that observed against the same variant in our Syrian hamster model (where the endpoint is at 4 days post infection) (19, 20). An improvement in lung pathology scores was also observed in the Molnupiravir-and Nirmatrelvir-treated SCID mice as compared to the vehicle-treated mice. Thus the SCID mice/beta variant infection model may serve as a useful tool to assess the *in vivo* efficacy of antiviral molecules against SARS-CoV-2.

It is surprising that infected SCID mice seem to control the infection by day 4 post infection. Moreover, monitoring a group of infected mice up to 14 days pi did not reveal any morbidity signs or weight loss over time. Typically infection of SCID mice with viruses (that are able to replicate in mice) results in a lethal infection (21–23).

The advantages of using this mouse model for initial *in vivo* evaluation of antivirals include; (i) the use of a real clinical isolate without the need to use mouse-adapted strains or genetically modified animals (ii) roughly 6-fold less of the test drug needed for the *in vivo* efficacy studies (average weight of a hamster is 120 gram versus 20 gram for mice), which will save a lot of material which is in particular important in case of highly priced or not easy to synthesize compounds, (ii) more convenient housing conditions since up to 5 mice can be co-housed in one cage versus 2 hamsters per cage, which is important for the capacity of the high biosafety animal facility. Consequently, such a model will enable testing more compounds in shorter period of time. On the other hand, the limitation of this model, is that unlike for hamsters, mice are only susceptible to the beta variant. Since small molecule inhibitors should have equipotent activity against all variants this is not of concern for studies with such drugs. However, for testing of therapeutic antibodies, infection models (in hamsters) with the different VoC will still be needed. Likewise, for vaccine studies, fully immunocompetent animals are needed, hence SCID mice are not useful for this purpose. Therefore, this SCID mice/beta variant infection model will be mainly advantageous for the evaluation of small molecule inhibitors of SARS-CoV-2 replication.

## Material and Methods Virus

The SARS-CoV-2 strain used in this study, the beta variant B.1.351 (hCoV-19/Belgium/rega-1920/2021; EPI_ISL_896474, 2021-01-11), was recovered from a nasopharyngeal swab taken from a patient with respiratory symptoms returning to Belgium in January 2021 (24). A passage two virus on Vero E6 cells was used for the study described here. Live virus-related work was conducted in the high-containment A3 and BSL3+ facilities of the KU Leuven Rega Institute (3CAPS) under licenses AMV 30112018 SBB 219 2018 0892 and AMV 23102017 SBB 219 20170589 according to institutional guidelines.

### Cells

Vero E6 cells (African green monkey kidney, ATCC CRL-1586) were cultured in minimal essential medium (Gibco) supplemented with 10% fetal bovine serum (Integro), 1% L-glutamine (Gibco) and 1% bicarbonate (Gibco). End-point titrations were performed with medium containing 2% fetal bovine serum instead of 10%.

### SARS-CoV-2 infection of SCID mice

In brief, 7-9 weeks old male severe combined immune deficient (SCID) mice were purchased from Janvier Laboratories. Mice were housed in individually ventilated cages with a maximum of five mice per cage. For infection, mice were anesthetized with isoflurane and inoculated intranasally with 40 µL containing 10^5^ TCID_50_ SARS-CoV-2 beta variant (day 0). At different time-points post-infection (pi), 10 animals were euthanized by intraperitoneal (IP) injection of 100 μL Dolethal (200 mg/mL sodium pentobarbital, Vétoquinol SA) for collection of lung tissues.

### Treatment Regimen

Male SCID mice were treated by oral gavage with either the vehicle (n=20) or Molnupiravir (EIDD-2801, n=14) at 200 mg/kg or Nirmatrelvir (PF-332, n=14) at 300 mg/kg, twice daily starting from D0, just before the infection with the Beta variant as described in the previous section. All treatments were continued for 3 consecutive days (thus until day 2 pi). Mice were monitored for appearance, behavior and weight. At day 3 pi, mice were euthanized. Lungs were collected and viral RNA and infectious virus were quantified by RT-qPCR and end-point virus titration, respectively. The left lungs were fixed in 4% formaldehyde for histopathological analysis.

### SARS-CoV-2 RT-qPCR

Lung tissues were collected after sacrifice and were homogenized using bead disruption (Precellys) in TRK lysis buffer (E.Z.N.A.^®^ Total RNA Kit, Omega Bio-tek) and centrifuged (10.000 rpm, 5 min) to pellet the cell debris. RNA was extracted according to the manufacturer’s instructions. RT-qPCR was performed on a LightCycler96 platform (Roche) using the iTaq Universal Probes One-Step RT-qPCR kit (BioRad) with N2 primers and probes targeting the nucleocapsid (25). Standards of SARS-CoV-2 cDNA (IDT) were used to express viral genome copies per mg tissue.

### End-point virus titrations

Lung tissues were homogenized using bead disruption (Precellys) in minimal essential medium and centrifuged (10,000 rpm, 5min, 4°C) to pellet the cell debris. To quantify infectious SARS-CoV-2 particles, endpoint titrations were performed on confluent Vero E6 cells in 96-well plates. Viral titers were calculated by the Reed and Muench method (2 6) using the Lindenbach calculator and were expressed as 50% tissue culture infectious dose (TCID_50_) per mg tissue.

### Histology

For histological examination, the lungs were fixed overnight in 4% formaldehyde and embedded in paraffin. Tissue sections (5 μm) were analyzed after staining with hematoxylin and eosin and scored blindly for lung damage by an expert pathologist. The scored parameters, to which a cumulative score of 1 to 3 was attributed, were the following: congestion, intra-alveolar hemorrhagic, apoptotic bodies in bronchus wall, necrotizing bronchiolitis, perivascular edema, bronchopneumonia, perivascular inflammation, peribronchial inflammation and vasculitis.

### Ethics

Housing conditions and experimental procedures were approved by the ethics committee of animal experimentation of KU Leuven (license P001/2021).

### Statistics

GraphPad Prism (GraphPad Software, Inc.) was used to perform statistical analysis. Statistical significance was determined using the non-parametric Mann Whitney U-test. P-values of <0.05 were considered significant.

## Acknowledgments

We thank Carolien De Keyzer, Lindsey Bervoets, Thibault Francken, Stijn Hendrickx, Niels Cremers for excellent technical assistance. We are grateful to Piet Maes for kindly providing the SARS-CoV-2 strain used in this study. We thank Prof. Jef Arnout and Dr. Annelies Sterckx (KU Leuven Faculty of Medicine, Biomedical Sciences Group Management) and Animalia and Biosafety Departments of KU Leuven for facilitating the animal studies. We thank the Histology department of KU Leuven for technical support for histopathological analyses. We also thank Fran Berlioz-Seux, Rob Jordan and Betsy Russell for helpful discussion.

## Author Contributions

R.A., C.S.F., S.J.F.K and J.N. designed the studies; R.A., S.J.F.K and R.B. performed studies. R.A. and B.W. analyzed data; R.A. made the graphs; B.W., D.J. and J.N. provided advice on the interpretation of data; R.A. and J.N. wrote the paper; S.D.J provided essential reagents; R.A., C.S.F., S.J.F.K and J.N. supervised the study; L.V., D.J. and J.N. acquired funding.

---

**Supplementary Fig. S1.**
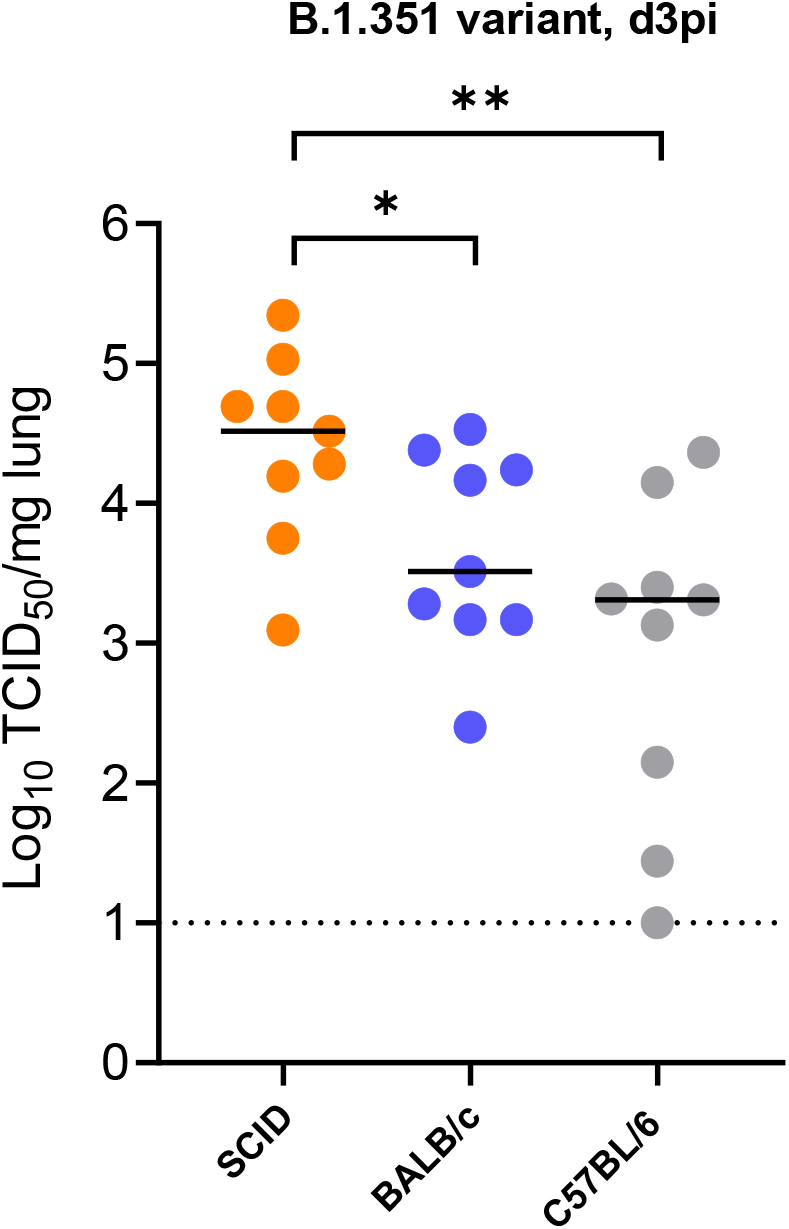
Replication of beta (B.1.351) SARS-CoV-2 in different mice. Infectious viral titers in the lungs of male SCID, male BALB/c and male C57BL/6 mice infected with 10^5^ TCID_50_ of beta SARS-CoV-2 variants at 3 days post-infection (pi) are expressed as log_10_ TCID_50_ per mg lung tissue. Individual data and median values are presented. Data were analyzed with the Mann™Whitney U test. *P < 0.05, **P<0.01). Data are from two independent experiment with n=9 per group.

## References

1. Adam D. 2022. 15 million people have died in the pandemic, WHO says. Nature. England https://doi.org/10.1038/d41586-022-01245-6.

2. Plante JA, Mitchell BM, Plante KS, Debbink K, Weaver SC, Menachery VD. 2021. The variant gambit: COVID-19’s next move. Cell Host Microbe 29:508–515.

3. Karim SSA, Karim QA. 2021. Omicron SARS-CoV-2 variant: a new chapter in the COVID-19 pandemic. Lancet 398:2126–2128.

4. Wan Y, Shang J, Graham R, Baric RS, Li F. 2020. Receptor Recognition by the Novel Coronavirus from Wuhan: an Analysis Based on Decade-Long Structural Studies of SARS Coronavirus. J Virol 94.

5. Liu Y, Hu G, Wang Y, Ren W, Zhao X, Ji F, Zhu Y, Feng F, Gong M, Ju X, Zhu Y, Cai X, Lan J, Guo J, Xie M, Dong L, Zhu Z, Na J, Wu J, Lan X, Xie Y, Wang X, Yuan Z, Zhang R, Ding Q. 2021. Functional and genetic analysis of viral receptor ACE2 orthologs reveals a broad potential host range of SARS-CoV-2. Proc Natl Acad Sci U S A 118.

6. Gu H, Chen Q, Yang G, He L, Fan H, Deng YQ, Wang Y, Teng Y, Zhao Z, Cui Y, Li Y, Li XF, Li J, Zhang NN, Yang X, Chen S, Guo Y, Zhao G, Wang X, Luo DY, Wang H, Yang X, Li Y, Han G, He Y, Zhou X, Geng S, Sheng X, Jiang S, Sun S, Qin CF, Zhou Y. 2020. Adaptation of SARS-CoV-2 in BALB/c mice for testing vaccine efficacy. Science (80-) 369.

7. Dinnon KH, Leist SR, Schäfer A, Edwards CE, Martinez DR, Montgomery SA, West A, Yount BL, Hou YJ, Adams LE, Gully KL, Brown AJ, Huang E, Bryant MD, Choong IC, Glenn JS, Gralinski LE, Sheahan TP, Baric RS. 2020. A mouse-adapted model of SARS-CoV-2 to test COVID-19 countermeasures. Nature 586:560–566.

8. Hassan AO, Case JB, Winkler ES, Thackray LB, Kafai NM, Bailey AL, McCune BT, Fox JM, Chen RE, Alsoussi WB, Turner JS, Schmitz AJ, Lei T, Shrihari S, Keeler SP, Fremont DH, Greco S, McCray PB, Perlman S, Holtzman MJ, Ellebedy AH, Diamond MS. 2020. A SARS-CoV-2 Infection Model in Mice Demonstrates Protection by Neutralizing Antibodies. Cell 182:744–753.e4.

9. Bao L, Deng W, Huang B, Gao H, Liu J, Ren L, Wei Q, Yu P, Xu Y, Qi F, Qu Y, Li F, Lv Q, Wang W, Xue J, Gong S, Liu M, Wang G, Wang S, Song Z, Zhao L, Liu P, Zhao L, Ye F, Wang H, Zhou W, Zhu N, Zhen W, Yu H, Zhang X, Guo L, Chen L, Wang C, Wang Y, Wang X, Xiao Y, Sun Q, Liu H, Zhu F, Ma C, Yan L, Yang M, Han J, Xu W, Tan W, Peng X, Jin Q, Wu G, Qin C. 2020. The pathogenicity of SARS-CoV-2 in hACE2 transgenic mice. Nature 583:830–833.

10. Liu FL, Wu K, Sun J, Duan Z, Quan X, Kuang J, Chu S, Pang W, Gao H, Xu L, Li YC, Zhang HL, Wang XH, Luo RH, Feng XL, Schöler HR, Chen X, Pei D, Wu G, Zheng YT, Chen J. 2021. Rapid generation of ACE2 humanized inbred mouse model for COVID-19 with tetraploid complementation. Natl Sci Rev 8.

11. Shuai H, Chan JFW, Yuen TTT, Yoon C, Hu JC, Wen L, Hu B, Yang D, Wang Y, Hou Y, Huang X, Chai Y, Chan CCS, Poon VKM, Lu L, Zhang RQ, Chan WM, Ip JD, Chu AWH, Hu YF, Cai JP, Chan KH, Zhou J, Sridhar S, Zhang BZ, Yuan S, Zhang AJ, Huang JD, To KKW, Yuen KY, Chu H. 2021. Emerging SARS-CoV-2 variants expand species tropism to murines. EBioMedicine 73.

12. Stolp B, Stern M, Ambiel I, Hofmann K, Morath K, Gallucci L, Cortese M, Bartenschlager R, Ruggieri A, Graw F, Rudelius M, Keppler OT, Fackler OT. 2022. SARS-CoV-2 variants of concern display enhanced intrinsic pathogenic properties and expanded organ tropism in mouse models. Cell Rep 38:110387.

13. Sun S, Gu H, Cao L, Chen Q, Ye Q, Yang G, Li RT, Fan H, Deng YQ, Song X, Qi Y, Li M, Lan J, Feng R, Guo Y, Zhu N, Qin S, Wang L, Zhang YF, Zhou C, Zhao L, Chen Y, Shen M, Cui Y, Yang X, Wang X, Tan W, Wang H, Wang X, Qin CF. 2021. Characterization and structural basis of a lethal mouse-adapted SARS-CoV-2. Nat Commun 12.

14. Halfmann PJ, Iida S, Iwatsuki-Horimoto K, Maemura T, Kiso M, Scheaffer SM, Darling TL, Joshi A, Loeber S, Singh G, Foster SL, Ying B, Case JB, Chong Z, Whitener B, Moliva J, Floyd K, Ujie M, Nakajima N, Ito M, Wright R, Uraki R, Warang P, Gagne M, Li R, Sakai-Tagawa Y, Liu Y, Larson D, Osorio JE, Hernandez-Ortiz JP, Henry AR, Ciouderis K, Florek KR, Patel M, Odle A, Wong L-YR, Bateman AC, Wang Z, Edara V-V, Chong Z, Franks J, Jeevan T, Fabrizio T, DeBeauchamp J, Kercher L, Seiler P, Gonzalez-Reiche AS, Sordillo EM, Chang LA, van Bakel H, Simon V, Alburquerque B, Alshammary H, Amoako AA, Aslam S, Banu R, Cognigni C, Espinoza-Moraga M, Farrugia K, van de Guchte A, Khalil Z, Laporte M, Mena I, Paniz-Mondolfi AE, Polanco J, Rooker A, Sominsky LA, Douek DC, Sullivan NJ, Thackray LB, Ueki H, Yamayoshi S, Imai M, Perlman S, Webby RJ, Seder RA, Suthar MS, García-Sastre A, Schotsaert M, Suzuki T, Boon ACM, Diamond MS, Kawaoka Y, group CMSPS (PSP) study. 2022. SARS-CoV-2 Omicron virus causes attenuated disease in mice and hamsters. Nature https://doi.org/10.1038/s41586-022-04441-6.

15. Chen Q, Huang X-Y, Liu Y, Sun M-X, Ji B, Zhou C, Chi H, Zhang R-R, Luo D, Tian Y, Li X-F, Hui Z, Qin C-F. 2022. Comparative characterization of SARS-CoV-2 variants of concern and mouse-adapted strains in mice. J Med Virol https://doi.org/10.1002/jmv.27735.

16. O’Toole Á, Hill V, Pybus OG, Watts A, Bogoch II, Khan K, Messina JP, Tegally H, Lessells RR, Giandhari J, Pillay S, Tumedi KA, Nyepetsi G, Kebabonye M, Matsheka M, Mine M, Tokajian S, Hassan H, Salloum T, Merhi G, Koweyes J, Geoghegan JL, de Ligt J, Ren X, Storey M, Freed NE, Pattabiraman C, Prasad P, Desai AS, Vasanthapuram R, Schulz TF, Steinbrück L, Stadler T, Parisi A, Bianco A, García de Viedma D, Buenestado-Serrano S, Borges V, Isidro J, Duarte S, Gomes JP, Zuckerman NS, Mandelboim M, Mor O, Seemann T, Arnott A, Draper J, Gall M, Rawlinson W, Deveson I, Schlebusch S, McMahon J, Leong L, Lim CK, Chironna M, Loconsole D, Bal A, Josset L, Holmes E, St George K, Lasek-Nesselquist E, Sikkema RS, Oude Munnink B, Koopmans M, Brytting M, Sudha Rani V, Pavani S, Smura T, Heim A, Kurkela S, Umair M, Salman M, Bartolini B, Rueca M, Drosten C, Wolff T, Silander O, Eggink D, Reusken C, Vennema H, Park A, Carrington C, Sahadeo N, Carr M, Gonzalez G, de Oliveira T, Faria N, Rambaut A, Kraemer MUG. 2021. Tracking the international spread of SARS-CoV-2 lineages B.1.1.7 and B.1.351/501Y-V2 with grinch. Wellcome open Res 6:121.

17. Owen DR, Allerton CMN, Anderson AS, Aschenbrenner L, Avery M, Berritt S, Boras B, Cardin RD, Carlo A, Coffman KJ, Dantonio A, Di L, Eng H, Ferre R, Gajiwala KS, Gibson SA, Greasley SE, Hurst BL, Kadar EP, Kalgutkar AS, Lee JC, Lee J, Liu W, Mason SW, Noell S, Novak JJ, Obach RS, Ogilvie K, Patel NC, Pettersson M, Rai DK, Reese MR, Sammons MF, Sathish JG, Singh RSP, Steppan CM, Stewart AE, Tuttle JB, Updyke L, Verhoest PR, Wei L, Yang Q, Zhu Y. 2021. An Oral SARS-CoV-2 Mpro Inhibitor Clinical Candidate for the Treatment of COVID-19. Science (80-) eabl4784.

18. Toots M, Yoon JJ, Cox RM, Hart M, Sticher ZM, Makhsous N, Plesker R, Barrena AH, Reddy PG, Mitchell DG, Shean RC, Bluemling GR, Kolykhalov AA, Greninger AL, Natchus MG, Painter GR, Plemper RK. 2019. Characterization of orally efficacious influenza drug with high resistance barrier in ferrets and human airway epithelia. Sci Transl Med 11.

19. Abdelnabi R, Foo CS, De Jonghe S, Maes P, Weynand B, Neyts J. 2021. Molnupiravir Inhibits Replication of the Emerging SARS-CoV-2 Variants of Concern in a Hamster Infection Model. J Infect Dis 224:749–753.

20. Abdelnabi R, Foo CS, Jochmans D, Vangeel L, De Jonghe S, Augustijns P, Mols R, Weynand B, Wattanakul T, Hoglund RM, Tarning J, Mowbray CE, Sjö P, Escudié F, Scandale I, Chatelain E, Neyts J. 2022. The oral protease inhibitor (PF-07321332) protects Syrian hamsters against infection with SARS-CoV-2 variants of concern. Nat Commun 13:719.

21. Neyts J, De Clercq E. 2001. Efficacy of 2-amino-7-(1,3-dihydroxy-2-propoxymethyl)purine for treatment of vaccinia virus (orthopoxvirus) infections in mice. Antimicrob Agents Chemother 45:84–87.

22. Charlier N, Leyssen P, Paeshuyse J, Drosten C, Schmitz H, Van Lommel A, De Clercq E, Neyts J. 2002. Infection of SCID mice with Montana Myotis leukoencephalitis virus as a model for flavivirus encephalitis. J Gen Virol https://doi.org/10.1099/0022-1317-83-8-1887.

23. Lefebvre DJ, De Vleeschauwer AR, Goris N, Kollanur D, Billiet A, Murao L, Neyts J, De Clercq K. 2014. Proof of Concept for the Inhibition of Foot-and-Mouth Disease Virus Replication by the Anti-Viral Drug 2’-C-Methylcytidine in Severe Combined Immunodeficient Mice. Transbound Emerg Dis 61:e89–e91.

24. Abdelnabi R, Boudewijns R, Foo CS, Seldeslachts L, Sanchez-Felipe L, Zhang X, Delang L, Maes P, Kaptein SJF, Weynand B, Vande Velde G, Neyts J, Dallmeier K. 2021. Comparing infectivity and virulence of emerging SARS-CoV-2 variants in Syrian hamsters. EBioMedicine 68:103403.

25. Boudewijns R, Thibaut HJ, Kaptein SJF, Li R, Vergote V, Seldeslachts L, Van Weyenbergh J, De Keyzer C, Bervoets L, Sharma S, Liesenborghs L, Ma J, Jansen S, Van Looveren D, Vercruysse T, Wang X, Jochmans D, Martens E, Roose K, De Vlieger D, Schepens B, Van Buyten T, Jacobs S, Liu Y, Martí-Carreras J, Vanmechelen B, Wawina-Bokalanga T, Delang L, Rocha-Pereira J, Coelmont L, Chiu W, Leyssen P, Heylen E, Schols D, Wang L, Close L, Matthijnssens J, Van Ranst M, Compernolle V, Schramm G, Van Laere K, Saelens X, Callewaert N, Opdenakker G, Maes P, Weynand B, Cawthorne C, Vande Velde G, Wang Z, Neyts J, Dallmeier K. 2020. STAT2 signaling restricts viral dissemination but drives severe pneumonia in SARS-CoV-2 infected hamsters. Nat Commun 11.

26. Reed LJ, Muench H. 1938. A simple method of estimating fifty per cent endpoints. Am J Epidemiol 27:493–497.

